# Quantifying the difference between CO_2_-release and carbon conversion in aerobic aquatic biodegradation tests

**DOI:** 10.1101/2025.03.20.644457

**Authors:** Ines Fritz, Lia Hergolitsch, Dennis Dalnodar, Antonia Zerobin

## Abstract

Quantitative biodegradation measurement of polymers (plastic) is, usually, done by monitoring indirect parameter, such as CH_4_- or CO_2_-release or oxygen consumption, which are a result of microbial activity. The microorganisms in such a test bottle will metabolise a polymer only if they can use a certain part of the substrate for growth of new biomass (new cells). This part can not be measured with the indirect parameter. In our long time study we investigated a wide range of fast to slowly biodegradable polymers by measuring the biomass growth in addition to the CO_2_-release. To that end we extended and improved the information about the metabolisation of the polymer for a full carbon balance. We quantified the biomass growth by measuring insoluble, therefore cellular, protein content from the inocula, from the suspensions during the biodegradation test runtime and at the end of each test. The procedure reduces the doubt about full biodegradability of all polymers, but especially that of mixtures and blends composed from different polymers. Our method evaluation is based on the results of 150 biodegradation tests.

## 1 Introduction

According to the European Union Directive (2019/904/EU) there is meanwhile no doubt about the need to replace conventional persistent plastic with biodegradable materials in all applications where whole products or parts of it can end up in the environment [3]. There is also no doubt about the fact that biodegradable plastic does neither reduce unintentional loss nor littering quantities – the expected effect is the reduced harm such losses cause to the ecosystems because of the finite lifetime of biodegradable materials [1].

However, there is some doubt in the public but also among scientists about ultimate biological conversion of polymers [12][20]. It derives from the necessity to measure the biodegradation progress by indirect parameters, such as CO_2_ evolution, biogas production, oxygen consumption or decay of total organic carbon (TOC) or dissolved organic carbon (DOC) in the test matrix [7]. Each of these methods has specific advantages and disadvantages, but all of them have in common that the conversion of polymer carbon into newly grown biomass carbon can not be measured [6].

The fate of a polymer‘s chemical elements can be monitored and quantified if it is synthesised from isotope marked components [16] or if the released CO_2_ reflects the known distribution of natural isotopes [22]. While such investigations are of basic scientific interest for studying the degradation and assimilation dynamics, they are not applicable for testing of materials and products deriving from a real-scale production facility and which are intended for certification (e.g. DIN-certco or TÜV Austria-Belgium „OK compost” label according to EN 13432) [4].

Textbooks about biochemistry and about microbiology have mentioned already in the early 20^th^ century the general principle of substrate utilisation by microorganisms [19]. One of these general principles is the universal bioconversion path from extracellular polymer cleavage, uptake of smaller molecules (monomers, oligomers), inside the cell then continuing with katabolism to generate universal small molecules (acetate) and eventually the division into anabolism and energy production [14]. Because of the generation of new biomass for growth and replication, not all of the incorporated carbon can and will become fully oxidised and released as CO_2_ (under aerobic conditions). Anaerobic metabolism may have different proportions between anabolism and energy generation but follows the same principal of life: a part of the substrate is used for growth.

**Figure 1.**
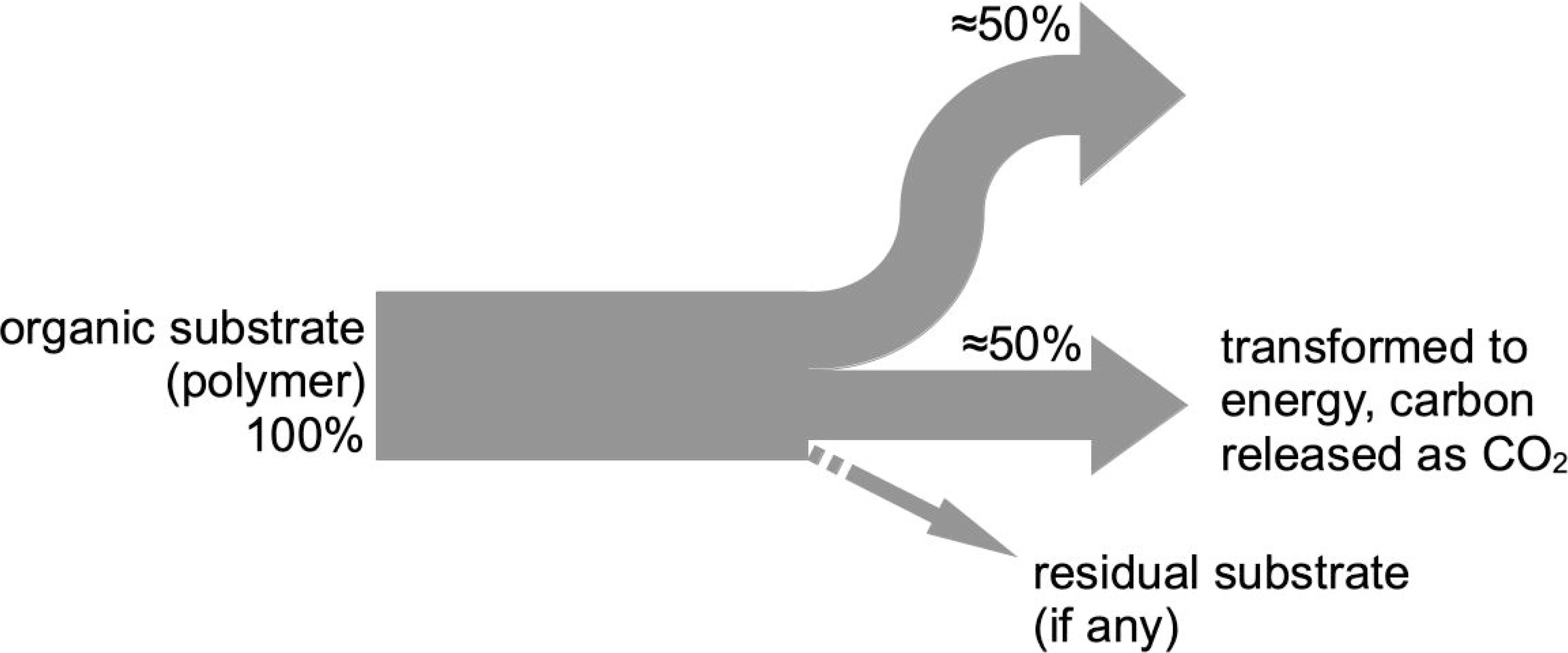
Carbon mass flow diagram: from substrate to end products due to microbial metabolisation (=biodegradation) [6].

It is therefore obvious that the above mentioned indirect measurements, as they are required for biodegradation testing, can never result in a complete release of the substrate carbon (e.g. all substrate carbon released as CO_2_) – measurement errors excluded. Scientists know about these principals of life and talk in casual speech about complete biodegradation of pure substances (such as polymers) when at least 60% of the substrate carbon is released as CO_2_. But the same scientists struggle with the interpretation of such CO_2_-release results when the test specimen was a mixture or blend of multiple substances (e.g. multiple polymers plus additives). Doubt about a parallel full biodegradation of each of the components seems justified, as diauxic growth prohibits that [19]. Furthermore, a minor component may not be biodegradable but may be overlooked because of its small contribution to the overall carbon balance. This all combined is the reason why the well known test criteria standards can at one hand not request 100% CO_2_ release rate but can not allow less than 90% as a pass level for assuring ultimate biodegradation.

The well known batch growth curve, as it is idealised shown in all textbooks, does not end with the stationary phase (the upper level plateau). Microbes can withstand starvation for a certain period of time but will eventually die out. When some bacteria cells disintegrate and release their content into the medium, others utilise this new substrate and start their metabolic activity again. Once more, the general path is followed, ending up in approx. 50-60% of the newly appeared carbon used for energy production and is released as CO_2_. This cycle can repeat over and over again until the last living cell consumes the remains of the second last – theoretically [19]. In praxis, each cycle lasts a bit longer than the one before (because of Michaelis-Menten kinetic [14]) and a minimum of three cycles of growth and decay have to happen until more than 90% of the initial substrate carbon converted into energy (and CO_2_) is reached – 55% with the first, 80% with the second, 91% with the third cycle. As a consequence, the 90% threshold level required by EN 13432 does not intend to allow 10% of the substrate polymer to remain undegraded, no, it requires significantly elongated test times for safety reasons (passing these three cycles of microbial growth and decay).

Even the approx. 50-60% carbon used for energy production is not a constant. The rate between anabolism and energy production is directly coupled with the required amount of maintenance energy [18]. In a summary, the less favourable the environmental conditions are for the biocenosis, the more energy is required to withstand the external factors and the more of the substrate carbon goes into energy production [9].

As mentioned before, only two situations allow a direct and quantitative measurement of the metabolic carbon pathways: (1) monitoring of isotope distributions from labeled substances or (2) measuring dissolved organic carbon in the test medium in case fully water soluble substances are investigated. For the usual routine biodegradation analysis of biodegradable plastic, none of these situations is applicable.

The draft standard ISO DIS 18957:2025 suggests to treat the solid fraction at the end of a test by dissolving the residual polymer material in organic solvents of choice and declare the insoluble remains as (newly grown) biomass [10]. Such a procedure can not be successful, as (1) parts of the biomass can be extracted with organic solvents and (2) typical polymer blends may not dissolve completely in one solvent. Using an extended series of solvent extractions under harsh conditions may completely destroy the biomass.

The draft standard ISO CD 23292:2025 suggests to quantitatively determine the newly grown biomass in a biodegradation test indirectly by the total DNA amount analysed via qPCR [11]. While this sounds reliable, the longer time presence of DNA which was released from dead cells must be considered [15][23]. Therefore, the measurement can be specific for the number of DNA strands, but is not necessarily reflecting the cellular mass, including a potentially variable amount of cellular storage substances [13].

There is a more promising option, which is the starting point of this study: Because of inability to separate living cells from residual insoluble polymer, another cellular component that is somehow correlating with the total cell mass has to be used. We followed the guideline of a singular publication that considers all those above mentioned facts and choose protein as indicating parameter [21]. By comparing the biomass at begin and end of a test and calculating the biomass growth, the amount of biodegraded but not as CO_2_ released polymer carbon can be estimated [5].

Having measured a proportion between the average total organic carbon and the total protein content of an inoculum should, theoretically, allow to calculate a proportion factor. This number should be specific for a certain inoculum as long as neither the environmental factors nor the metabolic activity do change. However, collecting the microbial biocenosis from a complex real-life ecosystem is not possible without also collecting a certain amount of habitat (water, minerals and other organic substances). Separating the microorganisms cells from other content is therefore a critical necessity. But even more, adapting the native population to the new laboratory conditions (without adding anything, especially no nutrients and no substrate) is required. By comparing the TOC to protein nitrogen proportion of the inoculum with the situation in the test blinds at the end of a test (which means several weeks of starvation) provides an additional information about the population composition after prolonged starvation – after going through multiple cycles of growth and decay.

Measuring the protein content of test vessels, which have gotten polymers or materials added, the grown biomass can be calculated by using the inoculum specific conversion factor. Adding the carbon content of the new biomass to the carbon which was released as CO_2_ should directly give the biodegraded (=metabolised) amount of the polymer or material. This can be expressed as percent of the carbon which was added to the test and represents biodegradation much better.

In this study we combine the results obtained during 27 years of analysis from 16 biodegradation experiments with a total of 62 investigated substances analysed from 150 biodegradation bottles to prove or disprove the possibility and to evaluate advantages and limits of calculating a carbon balance in aerobic aquatic biodegradation tests.

## 2 Methods

### 2.1 Polymer element analysis

The elements C, H, N and S were determined on a VARIO EL elemental analyser (Heraeus). A 5 mg sample together with WO_3_ catalyst (approx. 5 mg) was weighed into tin boats and carefully folded. The oxidative substance digestion was carried out at a combustion tube temperature of 1150 °C by explosive combustion of the sample in a highly oxygen-enriched (4.9) helium (5.0) atmosphere. In a downstream reduction tube (copper chips, 850 °C), nitrogen oxides (NOx), sulphur oxides (SOx) and volatile halogen compounds were chemically bound and eliminated. The gas components were quantified after selective adsorption and desorption on heatable separation columns using a thermal conductivity measuring cell (WLD).

Sulfanilic acid (Merck 100684) was used for instrument calibration and determination of the respective daily factor. The contents of the respective elements are given in mass percent [% w/w]. The oxygen content is calculated from the difference of the sum of the contents of C, H, N, S and ash to 100 %.

### 2.2 Inoculum preparation from sewage sludge

Sewage sludge was collected from the local 20000 EWG communal wastewater treatment plant (WWTP) in Tulln, Austria. The suspension was harvested from the pipe where enriched biomass was pumped from the sedimentation basin towards the biogas plant. The sludge was still aerobic.

The sludge was stirred in a 600 mL open glass baker in the laboratory at constant temperature of 21°C with passive aeration for a minimum of 5 days, but mostly 7 days prior to use. While the sludge did not contain any ready biodegradable solids after its retention time of 21 days in the WWTP, the additional time in the lab was provided for climate adaptation and stabilisation.

pH-value and solid dry matter was determined to characterise the inoculum, in some cases respiration activity was measured. However, the measured parameters did not have any significant influence on the biodegradability of biodegradable or not biodegradable polymers.

### 2.3 Inoculum preparation from compost

Compost was obtained from a local semi-industrial compost operation plant near our Institute. Compost is produced there from mixed communal, agricultural and green waste with typical seasonal variations. Still active and warm compost (Rottegrad 2, as defined in [2]) was collected for thermophilic biodegradation test, while ripe compost (Rottegrad 4-5) was used for tests at mesophilic conditions.

In both cases, approx. 1 kg compost was immediately submerged in approx. 4 litre warm or cold tap water, depending on the intended use, and manually stirred for one hour. Coarse particles were removed by sieving the suspension successively through sieves with 10, 2, 0,5 and 0,22 mm mesh widths. This suspension was allowed to sediment for 30 minutes. After this time swimming parts were removed from the top and sand and sediments were left behind when pouring the suspension carefully into a glass baker. As such, the suspension was stirred at room temperature or at 45°C, depending on the intended use, for a minimum of 3 and up to 5 days with passive aeration. After that time a similar sedimentation step was done after which the biomass was enriched and collected by a gentle centrifugation at approx. 500xg for 20 minutes. The pellet was then re-suspended in mineral medium (Table 1) and equilibrated for another 5 to 7 days at the intended test temperature.

**Table 1:**
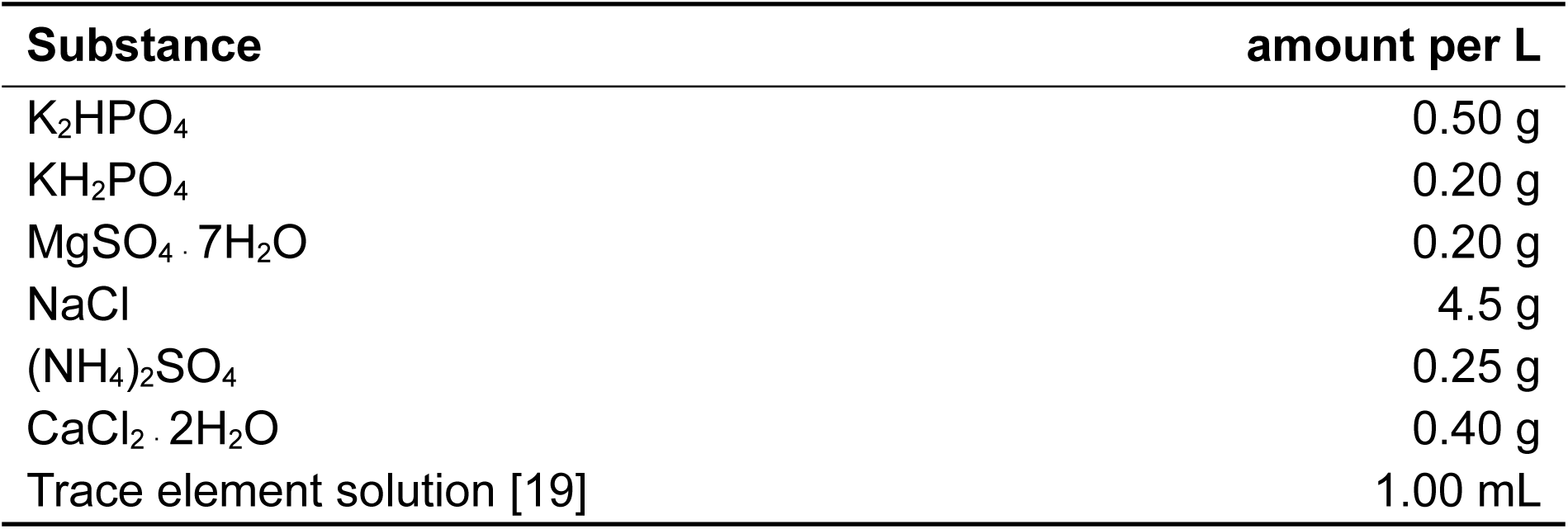
Mineral medium for the Sturm test, pH 7.1-7.2.

| Substance | amount per L |
| --- | --- |
| K<sub>2</sub>HPO<sub>4</sub> | 0.50 g |
| KH<sub>2</sub>PO<sub>4</sub> | 0.20 g |
| MgSO<sub>4</sub> · 7H<sub>2</sub>O | 0.20 g |
| NaCl | 4.5 g |
| (NH<sub>4</sub>)<sub>2</sub>SO<sub>4</sub> | 0.25 g |
| CaCl<sub>2</sub> · 2H<sub>2</sub>O | 0.40 g |
| Trace element solution [19] | 1.00 mL |

pH-value and solid dry matter was determined to characterise the inoculum, in some cases respiration activity was measured. However, the measured parameters did not have any significant influence on the biodegradability of biodegradable or not biodegradable polymers.

### 2.4 Aerobic aquatic biodegradation test (modified Sturm test)

Measurement of aerobic biodegradation via CO_2_-evolution was done following the procedure of OECD 301B, which is also called modified Sturm test [17]. We used 500 mL glass bottles (Schott) with gas washing heads. The test setup consisted of 400 mL carbon free mineral medium (Table 1), 10 mL of inoculum and an amount of test substance that contains roughly 500 mg organic carbon. The 250 mg ammonia sulfate equals approx. 53 mg nitrogen and allows for a synthesis of approx. 330 mg/L protein due to microbial growth.

The bottles were placed in an incubation chamber for thermophilic conditions (58 ± 2°C) or were operated in the laboratory at approx. 21 ± 1°C. Almost CO_2_-free air was pumped through the bottles in a rate of approx. 8-9 mL per minute and the exhaust gas was trapped in 0.2 M NaOH solution for each bottle individually. Absorbed CO_2_ was determined volumetrically by titration of an aliquot volume with 0.5 M HCl and calculated from the volume HCl between pH 9 (Phenolphthalein, Merck 107233) and pH 5 (mixed indicator Merck 106130). Titrations were done in intervals of two times per week at the begin and up to once every four weeks towards the end of a biodegradation test run.

When enough sample polymer was available, threefold determinations were done, but in some cases only double determination – and even then with reduced carbon content – was possible. Reference substances were either amorphous cellulose from spruce (Fluka 22181) or pure PHB (Sigma P-8150).

Evolved CO_2_ was expressed as percent of the theoretically possible CO_2_ calculated from the carbon content of the sample in each degradation bottle.

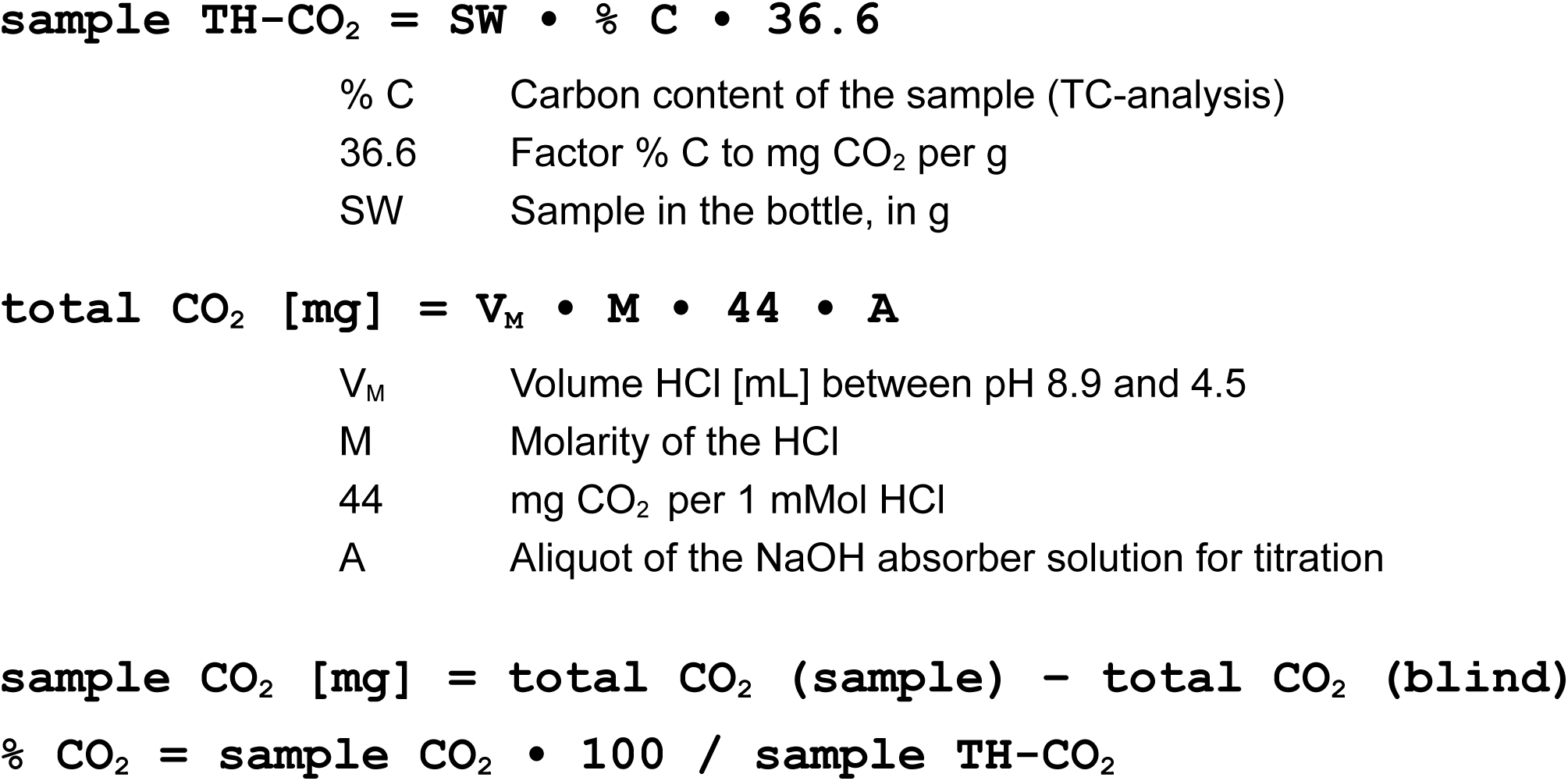

### 2.5 Determination of insoluble TC and TOC and soluble DC and DOC

Total carbon (TC) of suspensions (inocula, samples from degradation tests) was analysed in the Solid Sample Module (SSM-5000 A) of the TOC Analyser (5000 A, Shimadzu). Clear solutions (centrifugation supernatants, filtered through 0.22 µm PTFE) were directly injected into the TOC Analyser for measurement of dissolved carbon (DC). For analysis of the respective organic carbon, 2M HCl was added to the samples and CO_2_ gas was purged before thermal oxidation in the TOC Analyser – this was done by a procedure automatically run by the instrument.

TC and DC as well as TOC and DOC were analysed, TIC and DIC were calculated as the remains after subtracting the inorganic carbon part from the respective total carbon. The measurement limit for DOC and DIC was 2 mg/L.

After carefully examining the results from approx. 80 biodegradation tests (some not used as data base for this manuscript), we never found amounts of dissolved inorganic carbon above approx. 0.2% of the sample polymer carbon introduced to the test. Furthermore, we never found significant amounts of dissolved CO_2_ in inocula or in the blind tests at the end of the biodegradation tests. In those cases when we measured DC and DIC in samples taken while the biodegradation processes were still going on, we found DOC and DIC concentrations above the measurement limit, but which were still less than 1% of the carbon brought in with the polymer samples.

Some samples were analysed for TOC by using the ready-to-use test system of Hach-Lange (Art. Nr. LCK380) according to the producers procedure. The principle is similar to the instrumental determination in terms of organic substance digestion. The difference is in the photometric quantification of the released CO_2_ in an absorption tube adhered on top of the digestion tubet. The LCK380 test system is less sensitive than the instrumental analysis, therefore we never measured any concentration of DC or DIC above measurement limit (10 mg/L).

### 2.6 Determination of insoluble protein (total microbial cellular protein)

For the determination of biomass protein 10 mL aliquots from the degradation suspension were centrifuged (approx. 4500 g for 20 min), the supernatant was removed using a syringe with cannula and the pellet was washed once with de-ionised water, centrifuged again and pellets were stored frozen at -20°C until protein analysis.

400 µL of 0.25 M NaOH solution were added to each pellet and incubated in the water bath at 95°C for 15 minutes with gentle shaking, diluted by adding 600 µL deionised water and centrifuged at 4500xg for 5 minutes. 50 µL of the supernatant of the sample or a BSA standard was incubated with 1500 µL of Bradford reagent (Sigma-Aldrich No.: SLCG1879) for 30 minutes at room temperature. Measurement was then taken with a UV-Spectrophotometer (Shimadzu, UV-1800) at 595 nm. To calculate the protein concentration of the pellets, a calibration function with BSA standard in the concentration range between 0.005 mg/ml to 1 mg/ml was prepared.

### 2.7 Calculation of TC to protein relation and carbon balance

The carbon to nitrogen relation can be expressed based on different levels of organisation, but most often either as total organic carbon to nitrogen (TOC/N or more often simply C/N) or biomass to protein (BM/protein). Based on what we analysed with our methods we decided to not introduce empirical conversion factors, neither to calculate organic substance from total carbon nor calculate organic nitrogen from protein. While there are empirical factors for the C to N ratios of microbial cell mass given in literature (4.2 by [8], 3.57 by [14]), they differ quite a lot. It is to assume that differently composed inocula biocenoses may have differently composed cellular proteins, at least different to a certain degree. We decided to calculate the relation directly as we measure the parameter and therefore the relation of total carbon to protein (and neither organic carbon / protein nor carbon / nitrogen) was used for all further calculations.

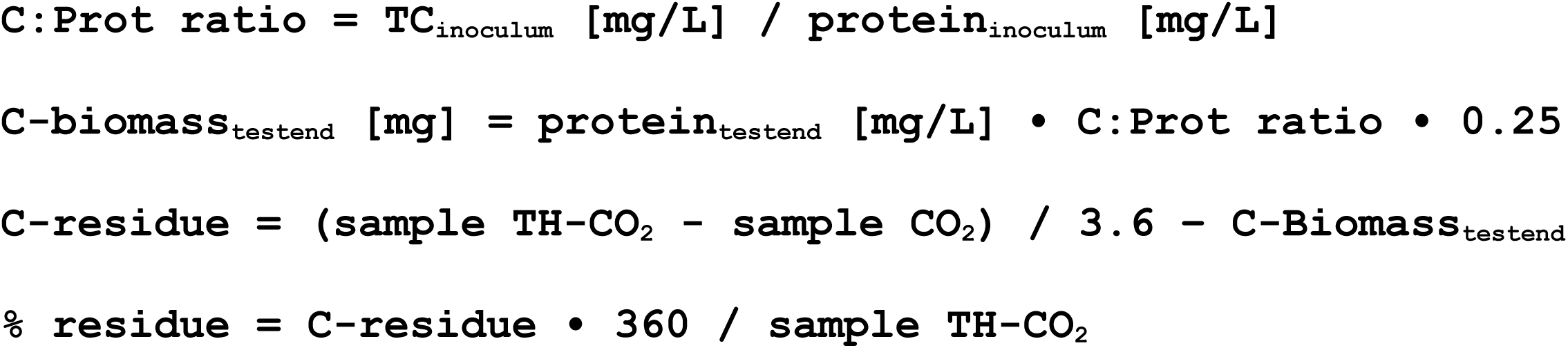

### 2.8 Statistics

All statistics (boxplots, outlier, error bars) and graphical representations were done with R-Studio (2024.12.1+563) with R version 4.4.3 (2025), using ggplot2, reshape2 and ggthemes libraries.

## 3 Results & Discussion

### 3.1 Biomass growth hypothesis test with water soluble substances

As described in the introduction, microbial metabolism prohibits a complete conversion of a substrate (not so for an inhibitory or toxic substance) into CO_2_, water and minerals. The driving force for substrate metabolisation is cellular growth – which is a increase of biomass. The usual aquatic biodegradation tests have to work with the assumption, that a release of approx. 50-60% of the substrate carbon as CO_2_ is equal to a full metabolic conversion of the substrates organic carbon.

For model experiments water soluble pure substances, namely starch (Sigma, S9765) and cellobiose (Sigma, C7252) were used. The first pre-experiment of this study was to measure CO_2_-evolution and substrate decay (as DC-decay) in an aquatic biodegradation test. The result is shown in Figure 2. Within two weeks both water soluble substrates were completely metabolised (97.6 % starch and 98.0 % cellobiose) but only 77 % of the substrates carbon was released as CO_2_ (77.0 % starch and 77.2 % cellobiose).

**Figure 2.**
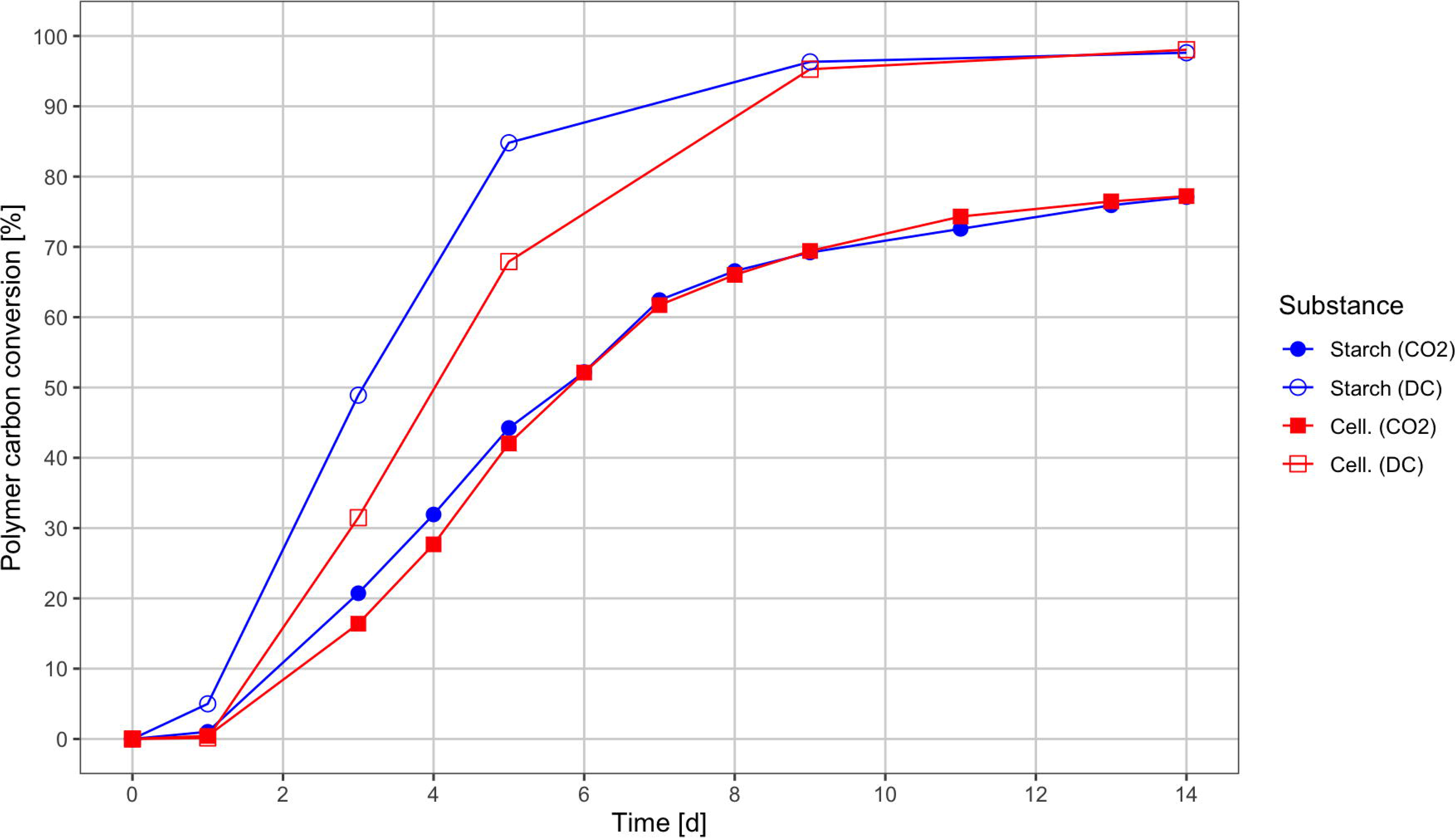
Comparison of CO_2_-evolution and DC-decay as measurement parameters in an aquatic biodegradation test. Mean values from three parallels, each.

CEN and ISO standard methods lack a description how to quantify the amount of organic carbon that derives from a substrate and ends up as new biomass (new living microorganisms cells). Biomass growth can be followed by usual microbiological and even by gravimetric methods in case a water soluble substrate is used. Testing insoluble polymers and blends of unknown composition disallows such simple biomass quantification measurements [6]. Other method principles have to be found, where the measurement of biomass must not be disturbed by residual particles of not biodegraded substrate. Therefore none of the more simple measurement methods, neither turbidity (OD) nor gravimetric, can be used and limitations of quantitative DNA analysis must be considered.

Protein that is released from dying cells may not be present in the medium for a longer time, as it is a highly valuable carbon-, energy- and nitrogen source for other cells. Active release of extracellular enzymes can accumulate to elevated concentrations during exponential growth but will be cleaved and absorbed in stationary and decay phase. Besides that, the protein content may not appear in a fixed relation to the total biomass of living cells over different activity phases. Biodegradation tests must be seen as batch cultivation with all the typical growth phases. During exponential growth high enzyme activity and initially low but rising contents of lipid and carbohydrate reserve may significantly change carbon / protein ratios over time. When activity is reduced during deceleration- and especially during stationary phase the carbon / protein relations later approaches a certain value asymptotically, which should eventually be similar to the initial value of an inoculum stabilized in the starvation state.

We therefore measured the time dependency of protein content in the aquatic biodegradation test with water soluble starch and cellobiose. When comparing the time dependent protein content in Figure 3 with the degradation progress in Figure 2, a direct correlation can be assumed. It is also seen, that the maximum protein content did not exceed the sum of initial concentration plus 330 mg possible from inorganic nitrogen of the medium. Biomass protein has increased from 95.7 mg/L at test begin to 220 and 189 mg/L for starch and cellobiose, respectively, within 14 days. The highest protein contents were reached at day 3 for starch (409 mg/L) and at day 5 for cellobiose (314 mg/L). Protein content of the blind (no substrate added) slowly decayed to 85.8 mg/L almost linearily.

**Figure 3.**
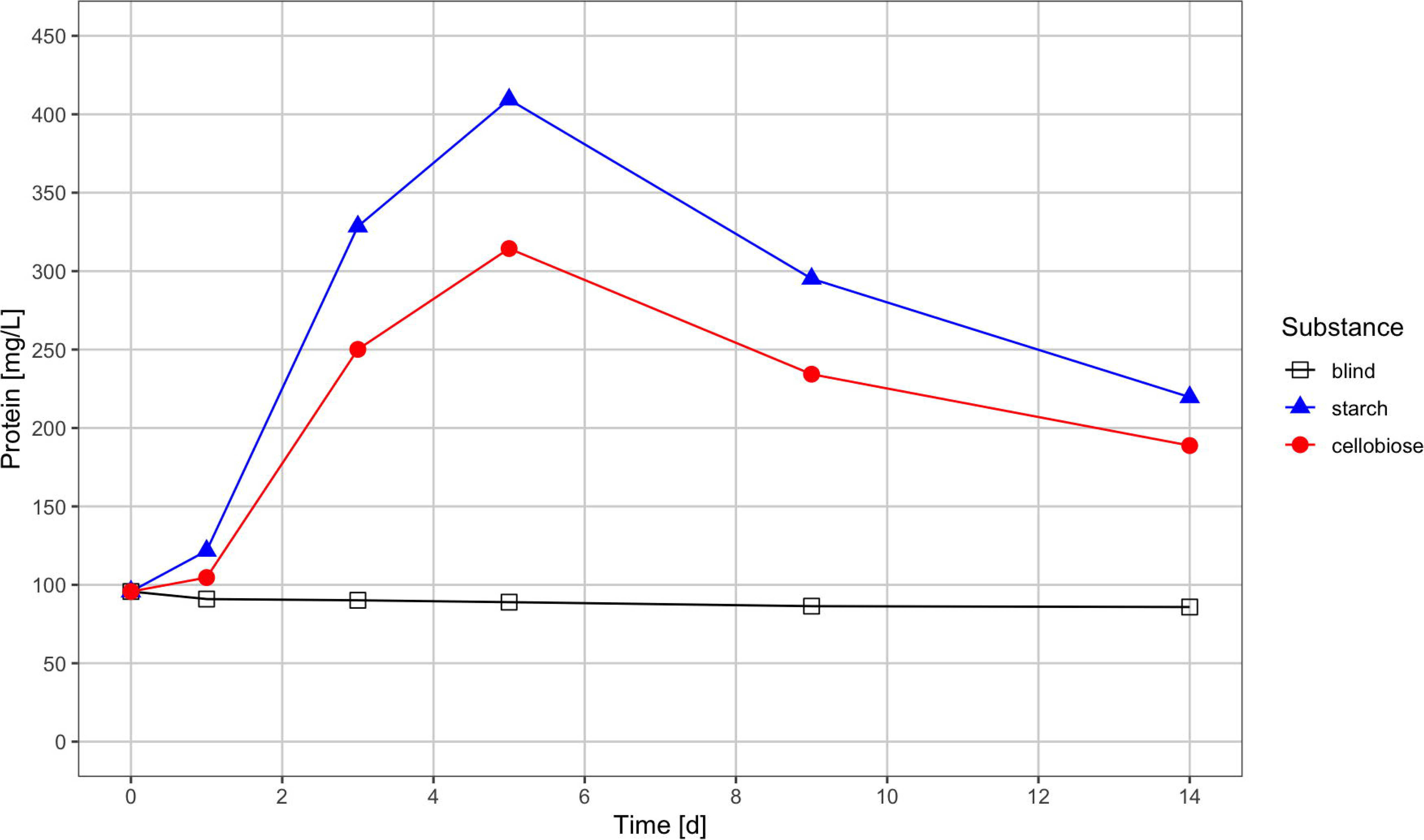
Protein content in the biodegradation bottles during starch and cellobiose degradation (same experiment as in Figure 2). Mean values from three parallels, each.

Measuring biomass organic carbon (the soluble fraction was removed by centrifugation) and biomass protein at different times from the same test finally showed the microbial metabolisation dynamic of the substrate. The carbon / protein relations are shown in Figure 4. Biomass carbon is given as total carbon (TC), no inorganic carbon was measured separately. Microorganisms grew due to metabolisation of the substrates according to their ease of digestibility, changed their carbon / protein relation, presumably because of increased protein synthesis, and shifted the relation back to higher numbers over time. Because of the short test time, the stationary phase was barely reached and the biocenosis in the substrate bottles had putatively different conditions compared to those of the blinds. The blinds held their carbon / protein relation almost constant – next to no biological activity was observed (17 mg CO_2_, equal to 4.6 mg C total released during these 14 days).

**Figure 4.**
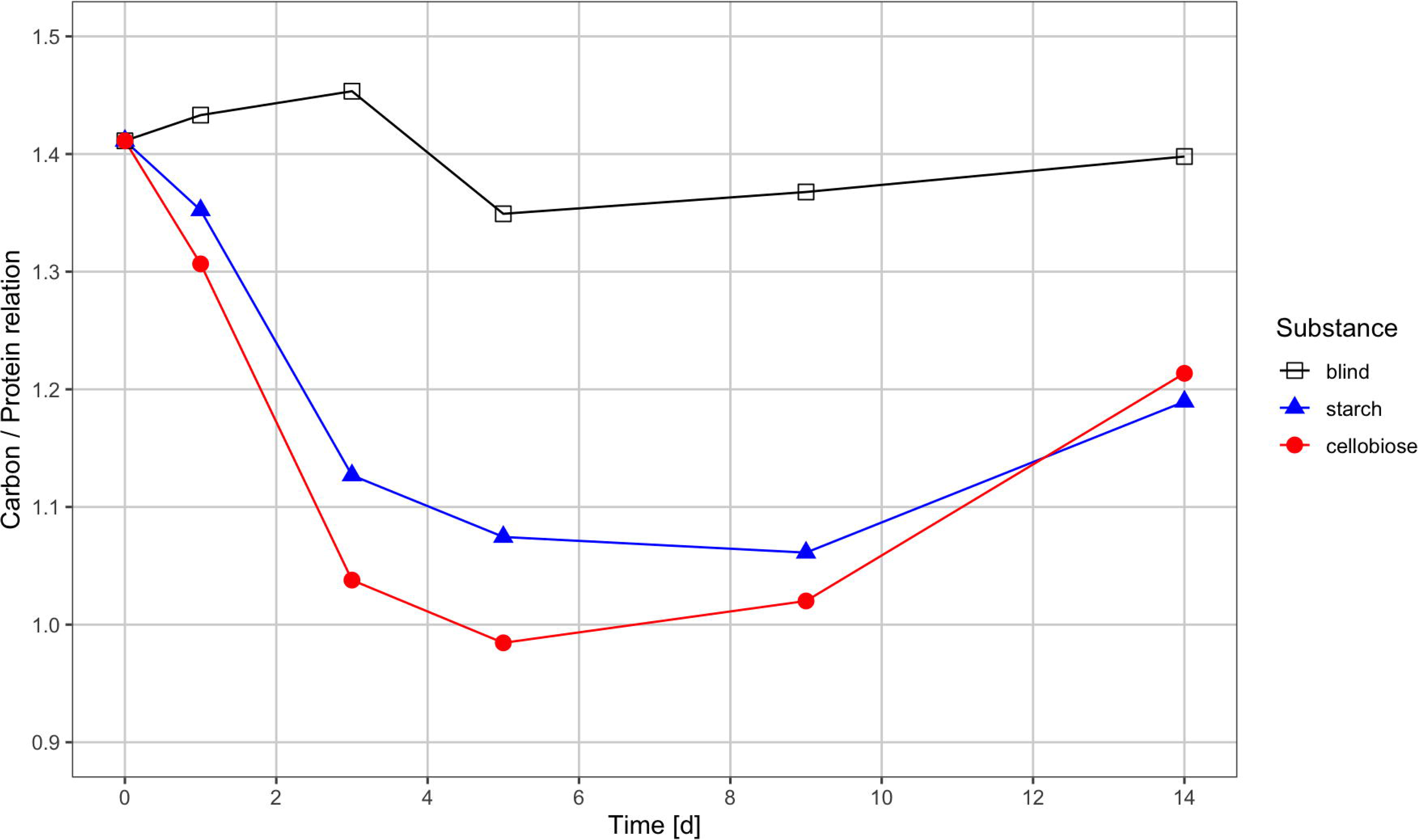
Relation of biomass carbon (TC) / biomass protein during a biodegradation test for a blind and for the two water soluble substances (same experiment as in Figure 2). Mean values from three parallels, each.

### 3.2 TC / protein relations of microbial inocula before and after a biodegradation test

Biomass carbon (expressed as TC) and biomass protein was measured for 18 inocula and 28 aquatic biodegradation test blinds. All samples were measured in double determination. At the end of each biodegradation test six carbon- and six protein results were obtained. Only two generally different inocula types were used, both in their native composition and without pretreatment or pre-exposure, but after removal of coarse particles (sand, swimming things) and after an equilibration time between 7 to 10 days at according temperature. The two types were: mixed microorganism suspensions from sewage sludge at the outflow of the active basin and washed-off suspensions from source separated communal biowaste compost.

The obtained carbon to protein relations of these inocula are shown in Figure 5. There was a significant difference in the initial numbers between sewage sludge and compost inocula. Both these inocula types contain living microorganisms and additional organics of unknown composition. Because of the equilibration time, we assume having used inocula that were already in or close to the stationary phase.

**Figure 5.**
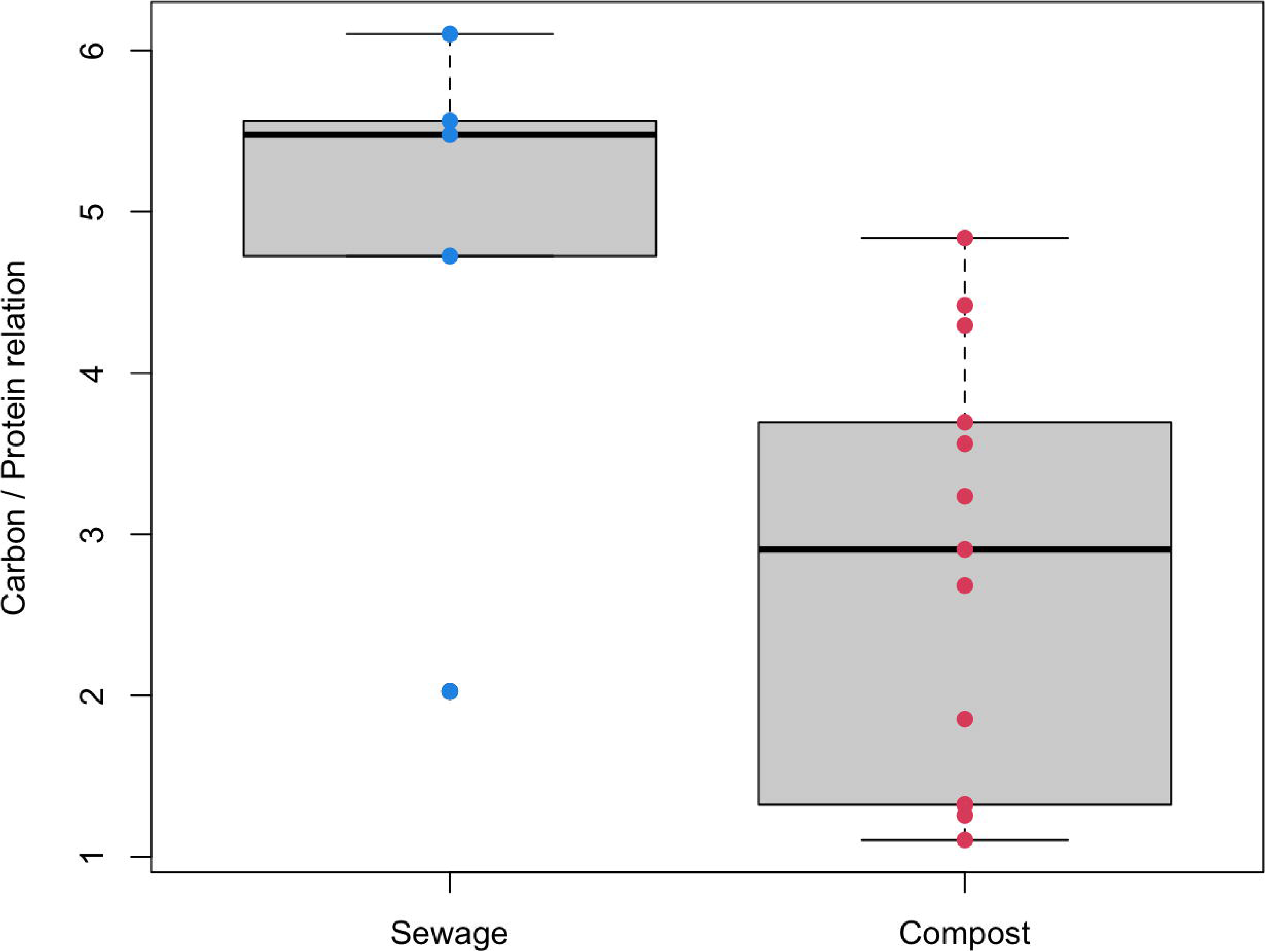
Boxplots of suspended organic carbon (as TC) / protein relations in different inocula. sewage sludge n = 5; compost n = 13

Total CO_2_-evolution from the blinds was generally low, with exceptions, indicating a low content of biodegradable organics which were brought into the test bottles with the inoculum. Bubbling the inflow air through 2 M NaOH did not remove all CO_2_ before entering the biodegradation bottles. Finally, it was not possible to distinguish between CO_2_-residues from the air inflow and CO_2_-evolution from microbial activity in the blinds. In one single experiment we used CO_2_-free synthetic air for aerating the biodegradation test (but did not do a full carbon balance analysis). In these tests we measured a CO_2_-release of 3.1 mg CO_2_ per g inoculum DM per month compared to a range of 3.6 to 5.7 mg CO_2_ per g inoculum DM per month in tests operated with compressed ambient air. Cost and effort of running aquatic biodegradation tests with synthetic air does not seem justified.

### 3.3 TC / protein relations after a biodegradation test

The biomass carbon to biomass protein relations of blinds at the end of the tests showed a wide range of values, graphically presented in Figure 6. Carbon to protein relations ranged from 1.2 up to 5.7. Blinds from thermophilic tests had a not significantly lower mean value of 2.8 compared to 3.2 in blinds from mesophilic tests. We can only assume that not controlled and not analysed factors had a bigger influence on the microbial biocenosis compositions than the test temperature had.

**Figure 6.**
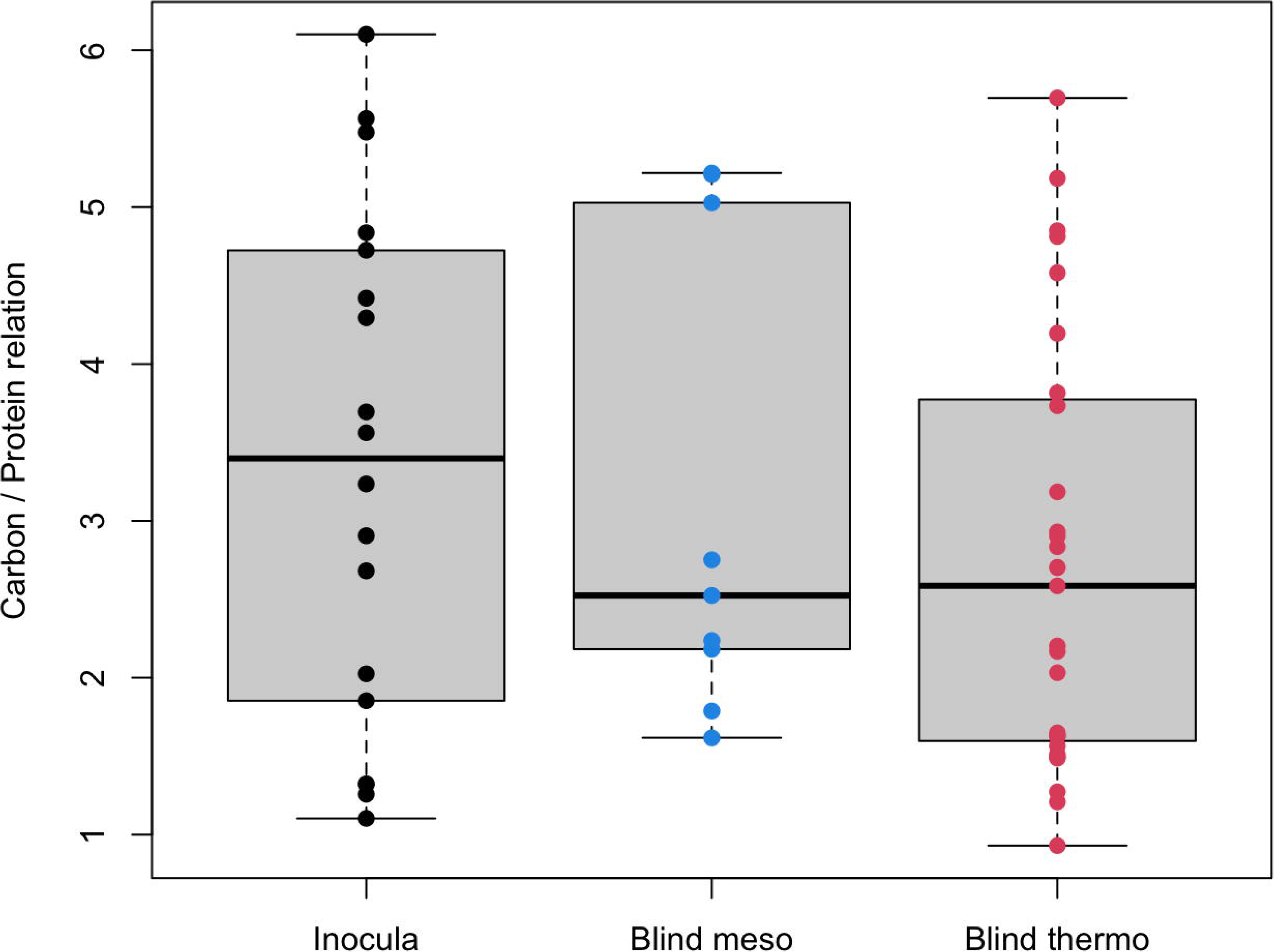
Boxplots of TC / protein relations from inocula and from the respective resulting biomass of blinds (no substrate added) after the according test duration. inokula n = 13; blind meso n = 9; blind thermo n = 19

### 3.4 Carbon balance method test with water soluble substances

After evaluating the prerequisites and specific analysis one-by-one, the generated data were combined into a full carbon balance calculation. By using water soluble substrates their disappearance during the biodegradation test could be followed analytically – which is impossible when not water soluble samples of, maybe, unknown composition were tested. Figure 7 shows the graphical representation of the special measurements for starch and Figure 8 shows the similar representation for cellobiose. For both these graphs the biomass carbon was not calculated from biomass protein measurement – the carbon content of the isolated biomass was directly determined.

**Figure 7.**
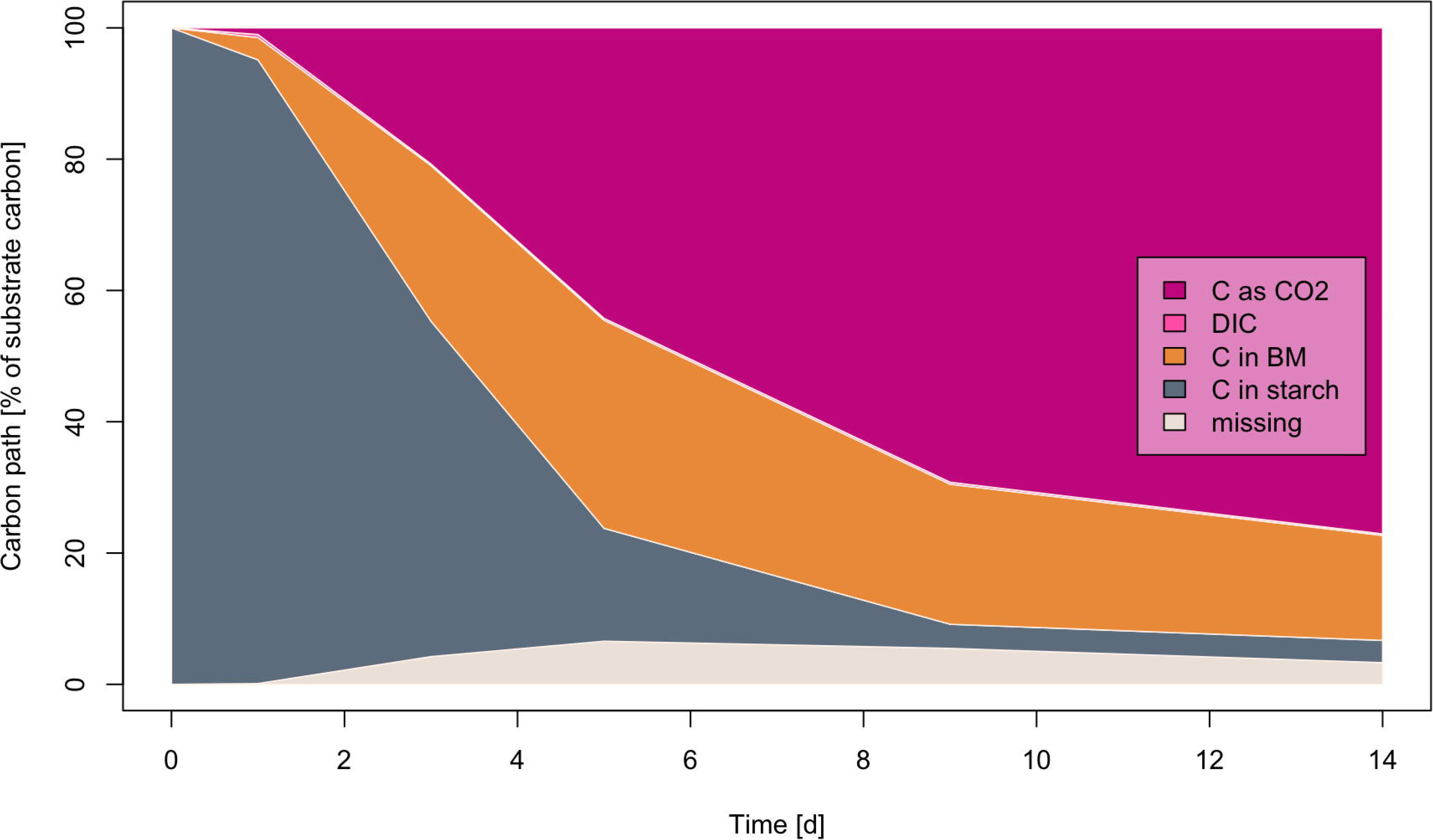
Biodegradation of water soluble starch expressed as CO_2_-evolution, dissolved CO_2_, biomass carbon and residuals (difference to 100%) for a full carbon balance (starch data as in Figure 2). Mean values from three parallels.

**Figure 8.**
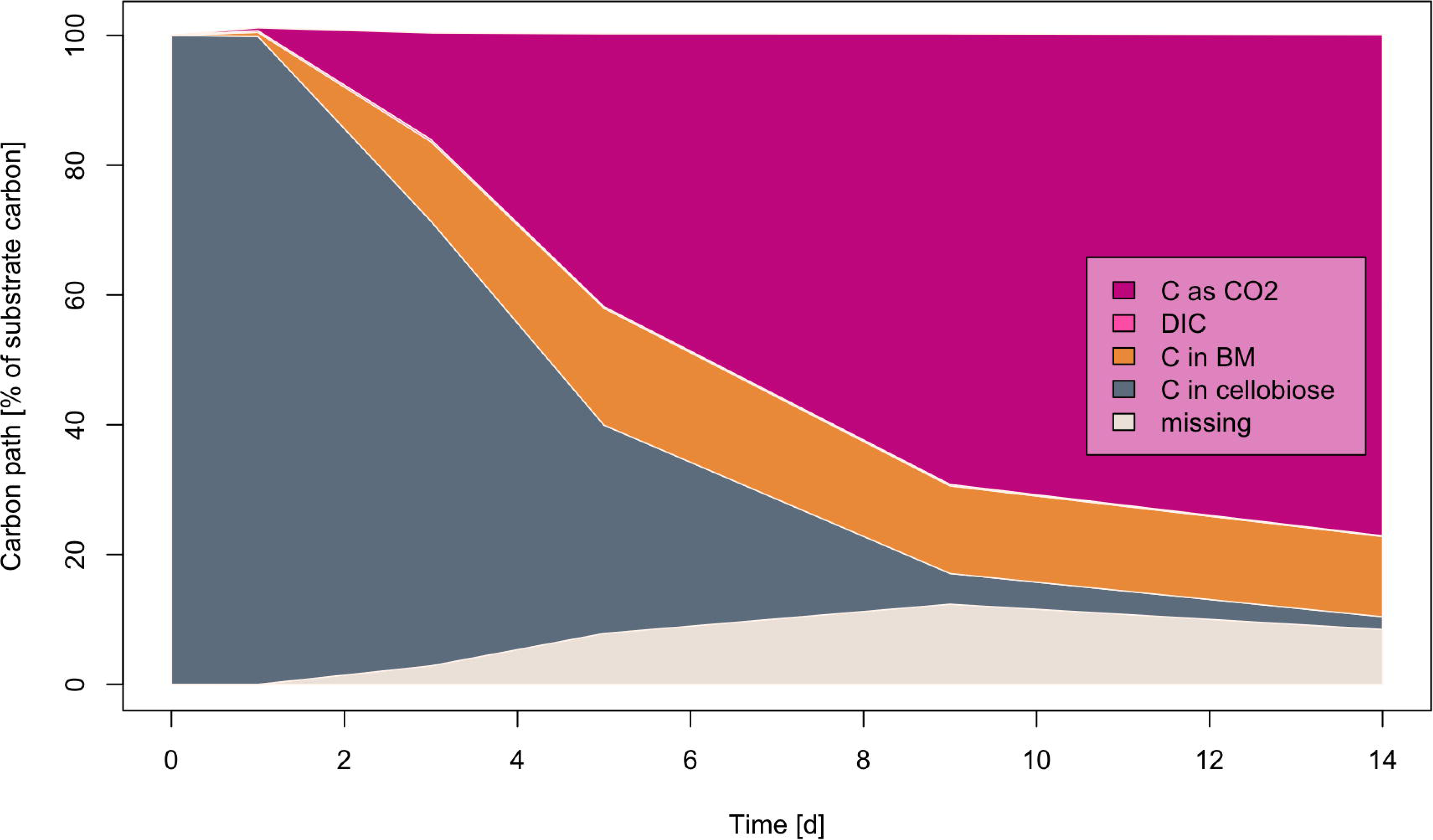
Biodegradation of water soluble cellobiose expressed as CO_2_-evolution, dissolved CO_2_, biomass carbon and residuals (difference to 100%) for a full carbon balance (cellobiose data as in Figure 2). Mean values from three parallels.

### 3.5 Method test with not water soluble, inherently biodegradable biopolymers

Water soluble substances allow the formulation of the hypothesis, but in practical routine analysis, next to all polymers are insoluble in water and have maybe unknown composition. There is no possibility to analytically determine if insoluble organic carbon derives from residual polymer or from newly grown biomass. A complete carbon balance, as shown before with the model substances, was not possible. Analysing the carbon to protein relation from the used inoculum is therefore a critical parameter, because the biomass carbon must be calculated from this relation, assuming that it doesn’t change much during the test time. The alternative – and therefore the control parameter – is the carbon to protein relation of the biomass obtained from the blinds at test end. However, due to prolonged starving conditions, compared to test bottles to which a biodegradable polymer was added, the relation may be different between polymer and blind bottles and different from the initial inoculum.

Results of an aquatic aerobic biodegradation test with starch and cellulose at mesophilic temperature (21°C) are summarised in Table 2. The carbon / protein relation of the inoculum was 5.38, and it was 3.00 in the blinds at the end of the test, after 36 days. While the indirect calculation of quantitative biodegradation via evolved CO_2_ would had resulted in 63% for cellulose and 68% for starch, carbon balance calculation increased these values to 92% and 77% degree of metabolisation for the two substances, respectively. The conspicuously high gap of almost 23% missing carbon for starch can be explained by a probably higher carbon to protein relation of the biomass grown on starch than that of the biomass grown on cellulose. However, due to the inability to measure the biomass carbon in these test suspensions directly, it remains a hypothesis. The comparably slower metabolised cellulose resulted in a good congruency between theory and praxis.

**Table 2.** Carbon balance for an aquatic aerobic biodegradation test with starch and cellulose at mesophilic conditions. Only two of the three parallels were analysed for carbon balance.

| Bottle | Polymer | Polymer C in [mg] | C as CO<sub>2</sub> out [mg]* | C in biomass [mg] | C as CO<sub>2</sub> out [%] | Metabolised [%] | Residue [%] |
| --- | --- | --- | --- | --- | --- | --- | --- |
| 2 | Cellulose | 413.8 | 257.0 | - | 62 | - | - |
| 4 | Cellulose | 479.0 | 322.7 | 116.1 | 67 | 92 | 8 |
| 8 | Cellulose | 367.6 | 219.4 | 118.7 | 60 | 92 | 8 |
| 3 | Starch | 469.7 | 336.1 | 40.1 | 72 | 80 | 20 |
| 5 | Starch | 363.6 | 248.7 | - | 68 | - | - |
| 6 | Starch | 429.7 | 270.9 | 50.1 | 63 | 75 | 25 |
| 1 | Blind | - | 82.1 | 8.9 | - | - | - |
| 7 | Blind | - | 75.6 | - | - | - | - |
| 9 | Blind | - | 80.7 | 9.5 | - | - | - |

\*) values given as net CO<sub>2</sub>-evolution from samples (individual sample minus mean value blinds)

### 3.6 Practical method check with a blend made from three biodegradable polymers

The distribution of substrate carbon (anabolism, energy production) is comparably well predictable when pure substances are investigated. It is much more challenging, if not impossible, to conclude from the amount of released CO_2_ to the degrees of metabolisation for each of the components of a mixed substrate (of a polymer blend). Therefore we tested the carbon balance procedure with a polymer blend consisting of three chemically different polymers, namely: 80%(w/w) PHBV (Enmat Y1000P, TianAn), 13.3%(w/w) PBS (Bio-PBS FD92PM, Mitsubishi), and 6.7%(w/w) PLA (Cordenka fibres 610F). Biodegradation is shown as diagram over time in Figure 9 and the data of the carbon balance are listed in Table 3. PHBV was used as positive control for this test due to limited amount of bottles.

**Figure 9.**
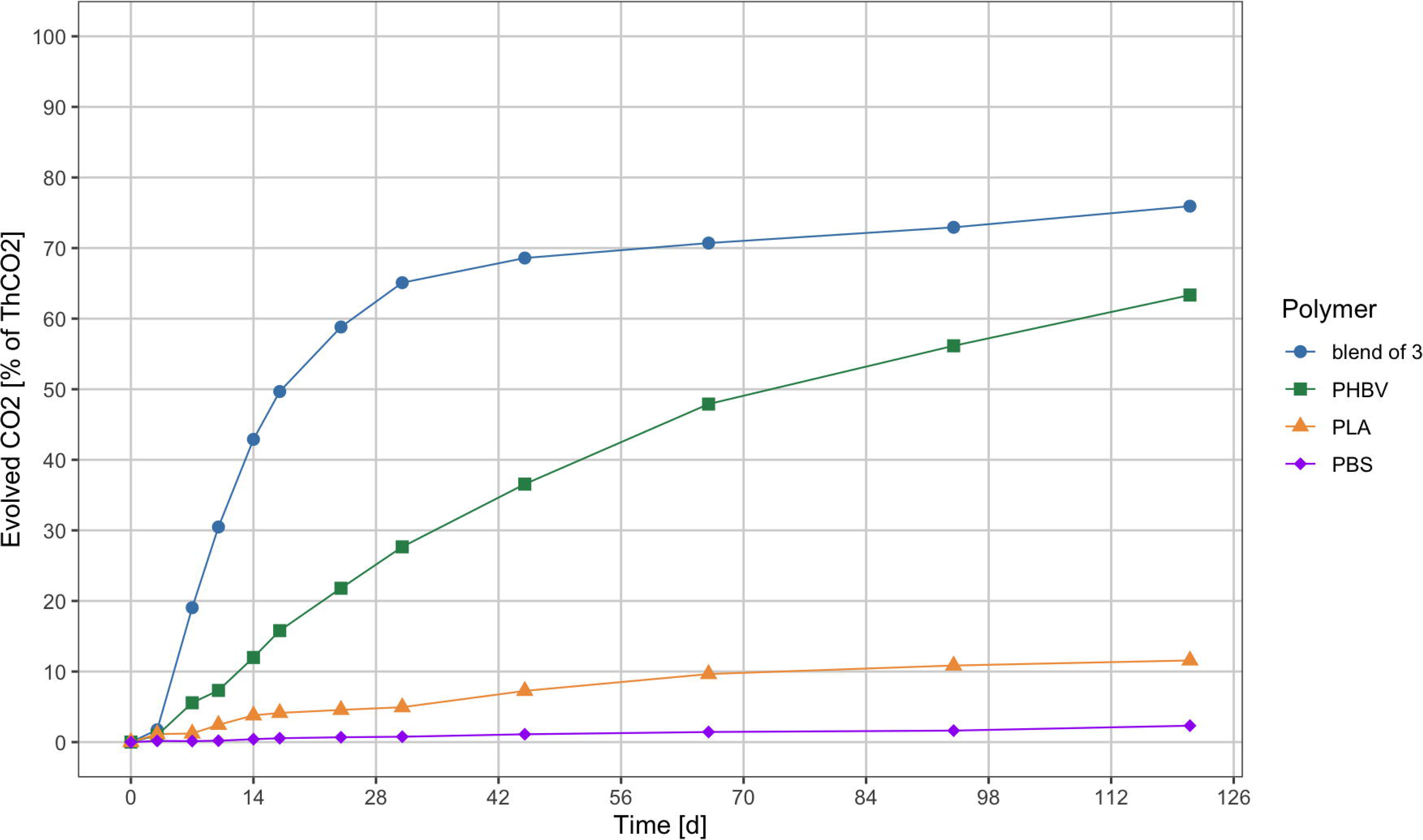
Biodegradation of PLA, PBS, PHBV and a blend of those three polymers in an aquatic aerobic biodegradation test at thermophilic conditions (58°C). Respective data in Table 2. All data are mean values from three parallels.

**Table 3.**
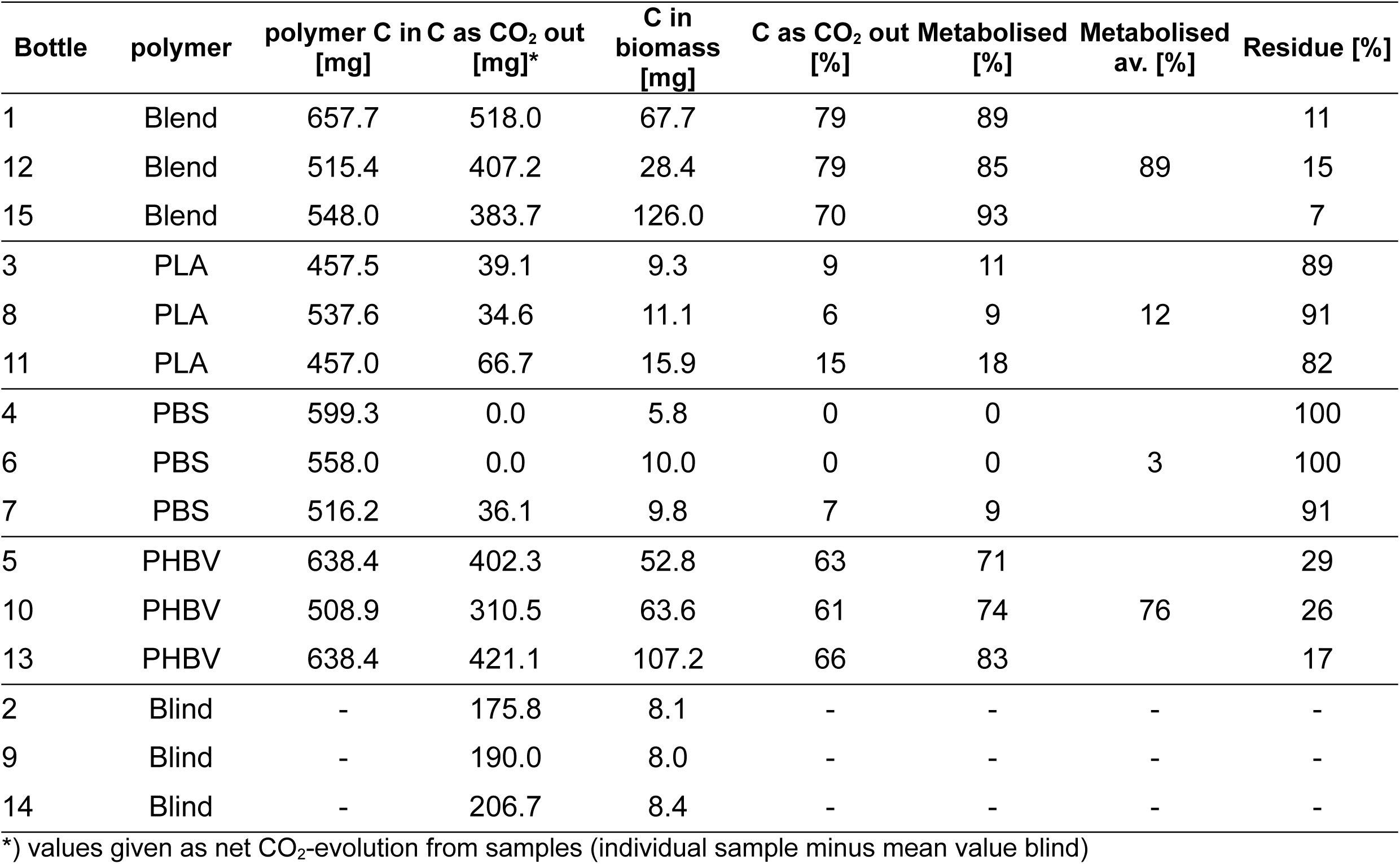
Carbon balance for an aquatic aerobic biodegradation test with a polymer blend consisting of three different biodegradable polymers at thermophilic conditions.

| Bottle | polymer | polymer C in [mg] | C as CO<sub>2</sub> out [mg]* | C in biomass [mg] | C as CO<sub>2</sub> out [%] | Metabolised [%] | Metabolised av. [%] | Residue [%] |
| --- | --- | --- | --- | --- | --- | --- | --- | --- |
| 1 | Blend | 657.7 | 518.0 | 67.7 | 79 | 89 |  | 11 |
| 12 | Blend | 515.4 | 407.2 | 28.4 | 79 | 85 | 89 | 15 |
| 15 | Blend | 548.0 | 383.7 | 126.0 | 70 | 93 |  | 7 |
| 3 | PLA | 457.5 | 39.1 | 9.3 | 9 | 11 |  | 89 |
| 8 | PLA | 537.6 | 34.6 | 11.1 | 6 | 9 | 12 | 91 |
| 11 | PLA | 457.0 | 66.7 | 15.9 | 15 | 18 |  | 82 |
| 4 | PBS | 599.3 | 0.0 | 5.8 | 0 | 0 |  | 100 |
| 6 | PBS | 558.0 | 0.0 | 10.0 | 0 | 0 | 3 | 100 |
| 7 | PBS | 516.2 | 36.1 | 9.8 | 7 | 9 |  | 91 |
| 5 | PHBV | 638.4 | 402.3 | 52.8 | 63 | 71 |  | 29 |
| 10 | PHBV | 508.9 | 310.5 | 63.6 | 61 | 74 | 76 | 26 |
| 13 | PHBV | 638.4 | 421.1 | 107.2 | 66 | 83 |  | 17 |
| 2 | Blind | - | 175.8 | 8.1 | - | - | - | - |
| 9 | Blind | - | 190.0 | 8.0 | - | - | - | - |
| 14 | Blind | - | 206.7 | 8.4 | - | - | - | - |

\*) values given as net CO<sub>2</sub>-evolution from samples (individual sample minus mean value blind)

PLA was slowly and PBS not at all metabolised within the 121 test days. PHBV was expected to biodegrade quickly at thermophilic conditions, the behaviour of the blend was unknown. Providing a biodegradation result solely based on the indirect measurement via evolved CO_2_ would had let questions open: does the approx. 76% conversion of the blends carbon into CO_2_ result from complete biodegradation of all it’s components? The carbon balance results in an amount of residue (11% by average) that is lower than the 13% of PBS the blend consists of. Therefore the carbon balance is able to show the effect of co-metabolism in the investigated blend and allows the conclusion of complete biodegradability within 121 days at thermophilic conditions.

### 3.7 Practical method test with biodegradation at different temperatures

PLA is well known to be slowly biodegradable at ambient/mesophilic conditions but is rapidly metabolised at temperature above approx. 55°C. We set up parallel aquatic aerobic tests at both temperatures and compared biodegradation and biomass growth for Cellulose and PLA. The combined biodegradation diagrams are shown in Figure 10 as an overlay and the carbon balance calculation data are listed in Table 4. Cellulose was fully metabolised at both temperatures, as expected. There was one outlier in the thermophilic test (bottle #8) which showed reduced CO_2_-evolution and reduced biomass growth. The reason for this deviation is unknown. Less than 3% of the PLA was metabolised during 62 days at mesophilic conditions, while 22% of the PLA was metabolised at thermophilic conditions during the same time. Biodegradation results based solely on CO_2_-evolution would had been approx. 30% lower in all cases. Calculating a carbon balance increased the confidence in biodegradation result and reduced doubts and or speculations about not-degraded residues.

**Figure 10.**
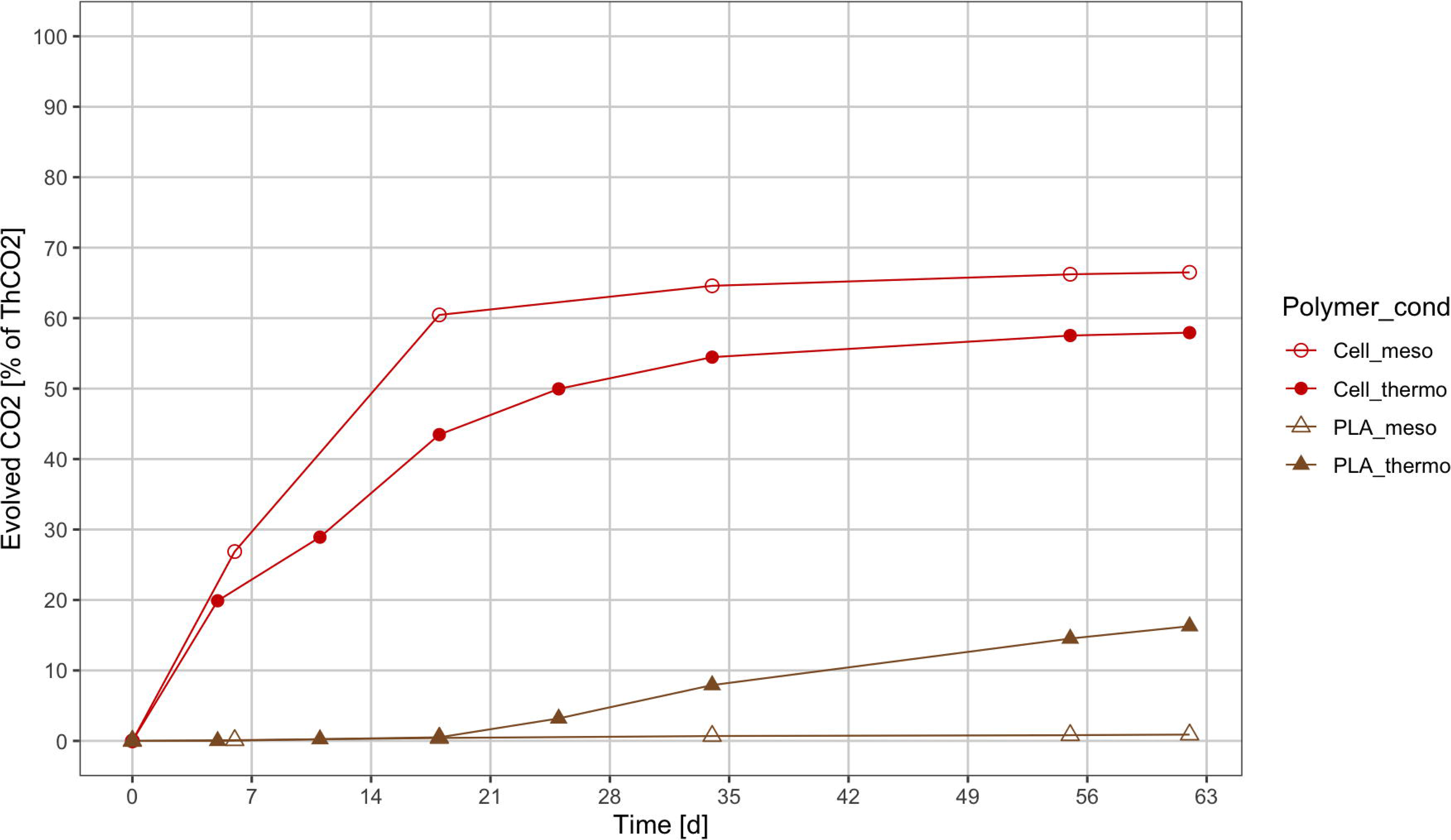
Biodegradation of PLA and cellulose via measurement of CO_2_-evolution at mesophilic (21°C) and thermophilic (58°C) conditions. All tests were done in triplicates.

**Table 4.** Carbon balance for two aquatic aerobic biodegradation tests with cellulose and PLA at mesophilic and thermophilic conditions. All data are average of triplicate analysis.

| mesophilic test |  |  |  |  |  |  |  |
| --- | --- | --- | --- | --- | --- | --- | --- |
| Bottle | Polymer | Polymer C in [mg] | C as CO<sub>2</sub> out [mg]* | C in biomass [mg] | C as CO<sub>2</sub> out [%] | Metabolised [%] | Missing [%] |
| 1 | Cellulose | 458 | 286 | 155 | 62 | 96 | 4 |
| 4 | Cellulose | 523 | 363 | 129 | 70 | 94 | 6 |
| 7 | cellulose | 494 | 333 | 121 | 67 | 92 | 8 |
| 5 | PLA | 690 | 3.5 | 11.6 | 1 | 2 | 98 |
| 6 | PLA | 786 | 10.2 | 12.2 | 1 | 3 | 97 |
| 9 | PLA | 686 | 0.0 | 11.7 | 0 | 2 | 98 |
| 2 | Blind | - | 79 | 10.6 | - | - | - |
| 3 | Blind | - | 89 | 13.3 | - | - | - |
| 8 | Blind | - | 75 | 12.3 | - | - | - |

| thermophilic test |  |  |  |  |  |  |  |
| --- | --- | --- | --- | --- | --- | --- | --- |
| Bottle | Polymer | Polymer C in [mg] | C as CO<sub>2</sub> out [mg]* | C in biomass [mg] | C as CO<sub>2</sub> out [%] | Metabolised [%] | Missing [%] |
| 5 | Cellulose | 576 | 365 | 188 | 63 | 96 | 4 |
| 6 | Cellulose | 532 | 317 | 198 | 60 | 97 | 3 |
| 8 | Cellulose | 551 | 280 | 158 | 51 | 79 | 21 |
| 1 | PLA | 625 | 131 | 44.8 | 21 | 28 | 72 |
| 3 | PLA | 717 | 95 | 36.6 | 13 | 18 | 82 |
| 9 | PLA | 626 | 91 | 40.1 | 15 | 21 | 79 |
| 2 | Blind | - | 109 | 13.3 | - | - | - |
| 4 | Blind | - | 106 | 14.8 | - | - | - |
| 7 | Blind | - | 96 | 16.8 | - | - | - |

\*) values given as net CO<sub>2</sub>-evolution from samples (individual sample minus mean value blind)

### 3.8 Practical method test from routine biodegradation analysis done within 27 years

Measurement data of 16 aquatic aerobic biodegradation tests with a total of 150 bottles were calculated for carbon balance and are provided in the support information <DATA in the Annex>. A wide range of natural and synthetic polymers, some of them in pure form, others as blends with known or partly unknown components were investigated. It doesn’t make sense to discuss each of those polymer biodegradation behaviours separately. Most interesting is, how much each of the quantitative biodegradation result was increased by adding newly grown microbial biomass carbon to the amount measured as evolved CO_2_. Over the 150 bottles the biodegradation results increased 17% by average and 22% for the cellulose references only. In only 12 of the 150 cases the sum built from evolved CO_2_ and biomass carbon exceeded 100% of the carbon introduced with the test substance, with the highest single value of 108%. According to these results it seems justified to claim that the information gain from doing a carbon balance exceeds by far the potential additional error deriving from uncertainties of the exact carbon to protein relation of the microbial biomass at a certain stage (at a certain day) of the biodegradation test. Carbon balance calculation should be included in European and International standard methods.

## 4 Conclusion

Routine biodegradation analysis of 62 polymer samples showed possibilities and limits of the aquatic tests method as well as allowed to evaluate the additional information gain from carbon balance analysis. The analysis of organic carbon and protein from biodegradation suspensions, which contain a mixture of residues from a water insoluble substrate (polymer) and microorganism cells of different life stages, is technically simple but has some pitfalls. Attention and care during biodegradation test operation and a general understanding of microbial dynamics is necessary for obtaining reliable carbon balance results.

### 4.1 Suggestions based on the data generated in this study

Minerals, floating things (wood, plastic, whatever) and excess suspended or dissolved organics should be removed from inocula. This can be achieved by sedimentation, sieving, gentle centrifugation (and careful separation of pellet and supernatant) and by equilibrating the suspension for several days at either laboratory environment or in the incubator at elevated temperature, according to the respective intention of meso- or thermophilic test conditions.

Carbon and protein content in the inoculum are critical parameters for calculating carbon balances. The initial relation of a fresh but stabilized inoculum reflects very well the biomass composition of the microorganisms at a later time of a test when the test substance turned out to be biodegradable. The inoculum parameters should be measured in three, better in five repetitions.

Measuring biomass carbon and biomass protein from the blinds of each test bottle at the end of the biodegradation test is essential for obtaining a reliable carbon balance for all those substances that are not or next to not biodegradable. The starvation situation of the blinds does not significantly differ from the starvation situation of a test bottle containing a not or next to not biodegradable substance. These measurements should be done with great care as well. Three parallel blind bottles are recommended and all parameters should be analysed in double determination from each of them separately. However, these analysis are less critical, as it doesn’t change the biodegradation result much when only some single percent of the substrate carbon was released as CO_2_ over many months.

Protein analysis is the base for calculating the biomass amount at test end and can be obtained by using the factor from either the inoculum or from the blinds at test end. The differences between those relations turned out to be not that big in a majority of tests but had exceptions in some, especially in tests operated at thermophilic conditions. All biodegradation tests should be done in triplicates and carbon balance should be calculated from each bottle individually. If the data are reliable, final values can be combined into average biomass growth, average CO_2_-release and average residual substrate carbon. Data can also be presented for each bottle individually and provided as ranges (from – to). Explanations for bigger deviations should be given.

Polymer blends and products containing raw protein as one of their components will disturb the protein based carbon balance calculation. We investigated one such product (data are property of a private company and not part of this study) and made a carbon balance after three month test time. The disturbance disappeared almost completely after this prolonged starvation time – excess organic nitrogen was slowly converted to nitrate by the microbial activity. It is important to get full information about the composition of a polymer before deciding on the test setup.

### 4.2 Remaining questions and suggested work for the future

The carbon to protein relation of a microbial test biocenosis seems to be within a certain range, but is not predictable, neither for the inoculum origin nor for the development during a test runtime for one given test situation. That is not a big surprise considering varying compositions and abundances of certain species depending on the substrate availability before harvest and depending on the chemical composition of the maybe unknown test specimen to be measured. No general rule, nor a general relation can be provided as of yet – each case is unique.

While we could demonstrate the general usefulness of measuring a carbon balance that justifies the additional effort in most of the cases, some mysteries remain, such as the gap differences shown in Table 2. In that case two known to be inherently biodegradable substances (starch and cellulose) were used as substrate with the more easily degradable showing a big amount of carbon missing in the combined analysis results. This should be taken as a hint about missing knowledge of metabolic reactions of a mixed microorganism community (in contrast to axenic single species cultures).

Despite presenting data from so many years of biodegradation measurement with 9 scientists and three technicians involved, the results were all obtained at one research institute and done with the varying instrumentations over time. Despite working with highest attention and care, we are looking forward to see other research groups repeating our methods and evaluating our data and findings. We would be pleased having suggested a valid method for calculating a carbon balance that is suitable for routine analysis and will, eventually, become the base for a recommendation in European and International standardisation.

## 6 Abbreviations used

BSA: bovine serum albumin (protein standard substance)
DOC: dissolved organic carbon
DC: dissolved carbon (organic and inorganic)
TOC: total organic carbon
TC: total carbon (organic and inorganic)
PCL: poly-caprolactone
PHB: poly-hydroxybutyrate
PHBV: poly-hydroxybutyrate-hydroxyvaleriate (biological random co-polymer)
PBS: poly-butyrate-succinate (synthetic co-polymer)
PLA: poly-lactic acid

## Annex – Data from biodegradation tests

<Annex 1 Data table.xlsx>

Carbon balance calculation for 62 polymer substances (including references) in 16 aquatic aerobic biodegradation tests. Explanations in the main text.

